# Diverse Changes in Microglia Morphology and Axonal Pathology Over One Year after Mild Traumatic Brain Injury in Pigs

**DOI:** 10.1101/2020.10.16.343103

**Authors:** Michael R. Grovola, Nicholas Paleologos, Daniel P. Brown, Nathan Tran, Kathryn L. Wofford, James P. Harris, Kevin D. Browne, John A. Wolf, D. Kacy Cullen, John E. Duda

## Abstract

Over 2.8 million people experience mild traumatic brain injury (TBI) in the United States each year, which may lead to long-term neurological dysfunction. The mechanical forces that occur due to TBI propagate through the brain to produce diffuse axonal injury (DAI) and trigger secondary neuroinflammatory cascades. The cascades may persist from acute to chronic time points after injury, altering the homeostasis of the brain. However, the relationship between the hallmark axonal pathology of diffuse TBI and potential changes in glial cell activation or morphology have not been established in a clinically relevant large animal model at chronic time points. In this study, we assessed tissue from pigs subjected to rapid head rotation in the coronal plane to generate mild TBI. Neuropathological assessments for axonal pathology, microglial morphological changes, and astrocyte reactivity were conducted in specimens out to 1 year post injury. We detected an increase in overall amyloid precursor protein pathology, as well as periventricular white matter and fimbria/fornix pathology after a single mild TBI. We did not detect changes in corpus callosum integrity or astrocyte reactivity. However, detailed microglial skeletal analysis revealed changes in morphology, most notably increases in the number of microglial branches, junctions, and endpoints. These subtle changes were most evident in periventricular white matter and certain hippocampal subfields, and were observed out to 1 year post injury in some cases. These ongoing morphological alterations suggest persistent change in neuroimmune homeostasis. Additional studies are needed to characterize the underlying molecular and neurophysiological alterations, as well as potential contributions to neurological deficits.

## Introduction

Traumatic brain injury (TBI) is a major health problem that is experienced by over 2.8 million people in the United States every year (48). Importantly, people who experience TBI often have permanent brain dysfunction and have a higher likelihood of developing chronic neurological disorders including major depressive disorder, post-traumatic stress disorder, and Alzheimer’s disease (19). TBI is typically caused by a mechanical insult that rapidly accelerates the head producing diffuse shear deformation forces throughout the brain. These mechanical forces can produce diffuse axonal injury (DAI), cytoskeletal fragmentation, synapse disruption, cellular membrane permeabilization, and blood brain barrier breakdown (8,11,23,41,47).

Subsequent secondary injury cascades may exacerbate initial brain trauma by amplifying astrocyte reactivity, leukocyte infiltration, and neuronal death which leads to tissue loss that can be more significant than the original insult (11,20,24,41). It is important to note that while glial cells play a central role in producing secondary injury, they also contribute to neuroprotection, plasticity, and regeneration. Indeed, recent studies found that microglia, the resident immune cell of the central nervous system, play a critical role in directing and facilitating tissue regeneration (10,15,31,37). Microglia also contribute to neuroinflammation by rapidly proliferating, secreting inflammatory cytokines, pruning axons and synapses, phagocytosing debris, and stabilizing the extracellular environment (9,18,27,36,38). Secondary injury cascades can persist in the brain for weeks, years, or even decades, as demonstrated in chronic assessments of mild TBI in rodents, acute assessments of mild TBI in pigs, and chronic assessments of moderate-to-severe TBI in humans (13,20,26–29). Moreover, the dynamic and diverse activity of microglia may also include region-specific responses to injury and tend to be more subtle following mild TBI (33,46).

In order to detect subtle morphological changes in microglia, sensitive and quantitative morphological analysis is needed. This type of analysis has not yet been applied to study the chronic effects of mild injury in pigs. Therefore, using a rotation-acceleration model of closed-head TBI in pigs, the current study seeks to histologically characterize the extent of axonal pathology and the resulting glial activity in the brain out to 1 year post-injury, with a particular focus on the diverse changes to microglia morphology. We hypothesized that specimens would display acute axonal pathology with morphologically altered microglia associating with regions of axonal pathology following a single mild TBI.

## Materials and methods

All procedures were approved by the Institutional Animal Care and Use Committees at the University of Pennsylvania and the Michael J. Crescenz Veterans Affairs Medical Center and adhered to the guidelines set forth in the NIH Public Health Service Policy on Humane Care and Use of Laboratory Animals (2015).

For the current study, specimens were obtained from a tissue archive of castrated male pigs subjected to a single mild TBI. This tissue archive was also used in Grovola et al (17). All pigs were 5 – 6 months-old, sexually mature (considered to be young adult), Yucatan miniature pigs at a mean weight of 34 ± 4 kg (total n = 29, mean ± SD). Pigs were fasted for 16 hours then anesthesia was induced with 20 mg/kg of ketamine and 0.5 mg/kg of midazolam. Following induction, 0.1 mg/kg of glycopyrrolate was subcutaneously administered and 50 mg/kg of acetaminophen was administered per rectum. All animals were intubated with an endotracheal tube and anesthesia was maintained with 2% isoflurane per 2 liters O_2_. Heart rate, respiratory rate, arterial oxygen saturation, and temperature were continuously monitored throughout the experiment. A forced-air temperature management system was used to maintain normothermia throughout the procedure.

In order to attain closed-head diffuse mild TBI in animals, we used a previously described model of head rotational-acceleration in pigs (7,42). Similar methods were described in Grovola et al (17). Briefly, each animal’s head was secured to a bite plate, which itself was attached to a pneumatic actuator and a custom assembly that converts linear motion into angular momentum. The pneumatic actuator rotated each animal’s head in the coronal plane, reaching an angular velocity between 165-185 radians per second (rad/sec) for the lower-level injured group (n = 4) and 230-270 rad/sec for the higher-level injured group (n = 15). Any presence of apnea was recorded, and animals were hemodynamically stabilized if necessary. Sham animals (n = 10) underwent identical protocols, including being secured to the bite plate, however the pneumatic actuator was not initiated. All animals were transported back to their housing facility, monitored acutely for 3 hours, and given access to food and water. Afterwards, animals were monitored daily for 3 days by veterinary staff.

At 3 days post-injury (DPI) (n = 4), 7 DPI (n = 4 at 165-185 rad/sec; n = 5 at 230-270 rad/sec), 30 DPI (n = 3), or 1 year post-injury (YPI) (n = 3), animals were induced and intubated as described above. Sham animals survived for 7 days (n = 4), 30 days (n = 1), or 1 year (n = 5). While under anesthesia, animals were transcardially perfused with 0.9% heparinized saline followed by 10% neutral buffered formalin (NBF). Animals were then decapitated, and tissue stored overnight in 10% NBF at 4°C. The following day, the brain was extracted and post-fixed in 10% NBF at 4°C for one week. To block the tissue, an initial coronal slice was made immediately rostral to the optic chiasm. The brain was then blocked into 5 mm thick coronal sections from that point by the same investigator. This allowed for consistent blocking and section coordinates across animals. All blocks of tissue were paraffin embedded and 8 μm thick sections were obtained using a rotary microtome.

Four sections from each specimen – one containing striatal tissue (approximately 10 mm anterior to the optic chiasm), one containing anterior aspects of hippocampal tissue (approximately 10 mm posterior to the optic chiasm), one containing posterior aspects of hippocampal tissue (approximately 15 mm posterior to the optic chiasm), and one containing cerebellar tissue (approximately 35 mm posterior to the optic chiasm) – were used for the ensuing Amyloid Precursor Protein (APP) histological analysis. Additional histological analysis examined only two sections from each specimen - one containing anterior aspects of hippocampal tissue and one containing posterior aspects of hippocampal tissue, as these sections displayed increased APP pathology in special neuroanatomical regions. Histological analysis of the corpus callosum only included sections with anterior hippocampal tissue, as sections with posterior hippocampal tissue did not contain corpus callosum.

For 3,3’-Diaminobenzidine (DAB) immunohistochemical labeling, we used a protocol outlined in Johnson et al (23). Briefly, slides were also dewaxed in xylene, rehydrated in ethanol and de-ionized water. Antigen retrieval was completed in Tris EDTA buffer pH 8.0 using a microwave pressure cooker then blocked with normal horse serum. Slides were incubated overnight at 4°C using either an anti-mouse APP (22C11) (Millipore, MAB348, 1:80,000), an anti-mouse GFAP (SMI-22) (Millipore, NE1015, 1:12,000), or an anti-rabbit Iba-1 (Wako, 019-19741, 1:4000) primary antibody. The following day, slides were rinsed in PBS and incubated in a horse anti-mouse/rabbit biotinylated IgG secondary antibody (VECTASTAIN Elite ABC Kit, Vector Labs, PK-6200). Sections were rinsed again, then incubated with an avidin/biotinylated enzyme complex (VECTASTAIN Elite ABC Kit), rinsed again, and incubated with the DAB enzyme substrate (Vector Labs, SK-4100) for 7 minutes. Sections were counterstained with hematoxylin, dehydrated in ethanol, cleared in xylene, and finally coverslipped using cytoseal. All sections were stained in the same histological sample run. All sections were imaged and analyzed at 20x optical zoom using an Aperio CS2 digital slide scanner (Leica Biosystems Inc., Buffalo Grove, IL).

For Luxol Fast Blue (LFB) staining, slides were dewaxed in xylene, and rehydrated in ethanol and deionized water. Slides were placed in a solution of 0.1% Solvent Blue 38 (Sigma, S-3382) and 95% ethanol warmed to 60°C for 4 hours, then differentiated in a lithium carbonate solution followed by 70% ethanol. Slides were counterstained in cresyl violet solution (Sigma, C5042), dehydrated in ethanol, cleared in xylene, and finally coverslipped using cytoseal. All slides were stained for LFB in the same histological sample run.

For APP semi-quantitative analysis, we initially characterized four specimens (three 7 DPI and one sham) and stained sections every 5 mm throughout the brain and brainstem for APP. Based on these slides, we identified six anatomical regions that contained APP pathology: periventricular white matter, striatum, ventral thalamus, dorsal thalamus, fimbria/fornix, and cerebellum. These regions were assessed by two blinded observers in the four previously described tissue sections for every specimen and given a 0-3 pathological burden score based on the amount of APP+ axons in the region (Figure 1a-c). The scores were summed then divided by the number of anatomical regions to provide a single, averaged pathological score for each specimen.

**Figure 1.**
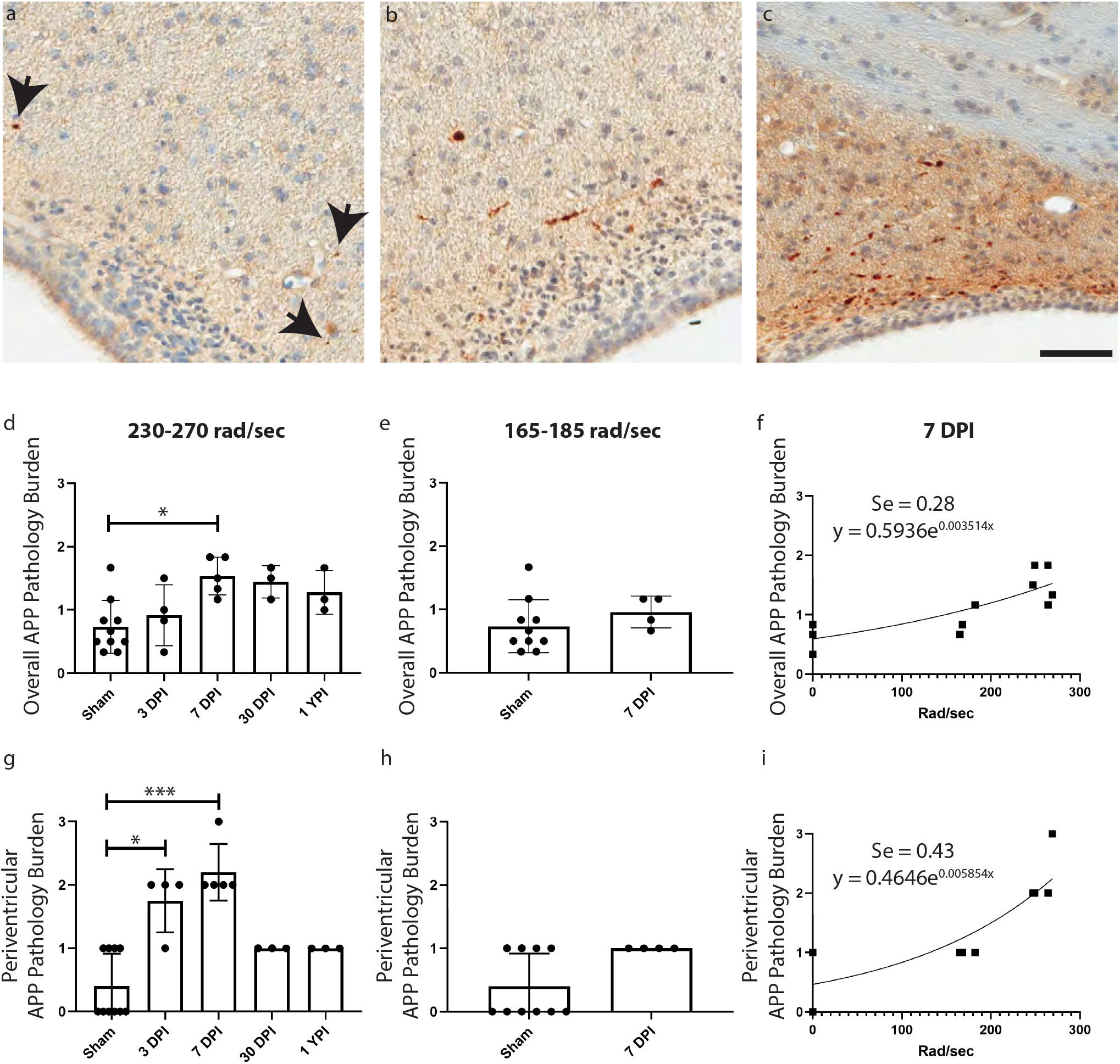
Examples are shown of APP pathology burden scored as a 1, 2, or 3 (a-c respectively, scale = 100 μm). The representative image in (a) is from a specimen injured at 165-185 rad/sec and survived to 7 days post injury (DPI), while images in (b) and (c) are from specimens injured at 230-270 rad/sec and survived to 7 DPI. The overall APP pathology burden for specimens injured at 230-270 rad/sec (d) and 165-185 rad/sec (e) are represented in the bar graphs. There is a significant change in APP pathology burden at 7 DPI in the 230-270 rad/sec group compared to sham (p = 0.0193). A non-linear regression line for the overall APP pathology burden of all 7 DPI specimens is shown (Se = 0.28) (f). Moreover, the APP pathology burden in the periventricular white matter increased at 3 DPI (p = 0.0372) and 7 DPI (p = 0.0008) compared to sham in the 230-270 rad/sec group (g), but did not change in the 165-185 rad/sec group (h). A non-linear regression line for the periventricular white matter APP pathology burden of all 7 DPI specimens is shown (Se = 0.43) (i).

For astrocyte semi-quantitative analysis, hippocampus and periventricular white matter were assessed in 2 sections per specimen, as well as inferior temporal gyrus and cingulate gyrus – two anatomical regions without APP pathology. We have adapted a semi-quantitative scale from Sofroniew et al. to histologically classify the progressive severity of reactive astrocytes (44). Each region was given a 0-3 glial fibrillary acidic protein (GFAP) reactivity score based on cell body size and density of GFAP+ cells in the region (Figure 2a-d). The scores were summed then divided by the number of anatomical regions to provide a single, averaged reactivity score for each specimen.

**Figure 2.**
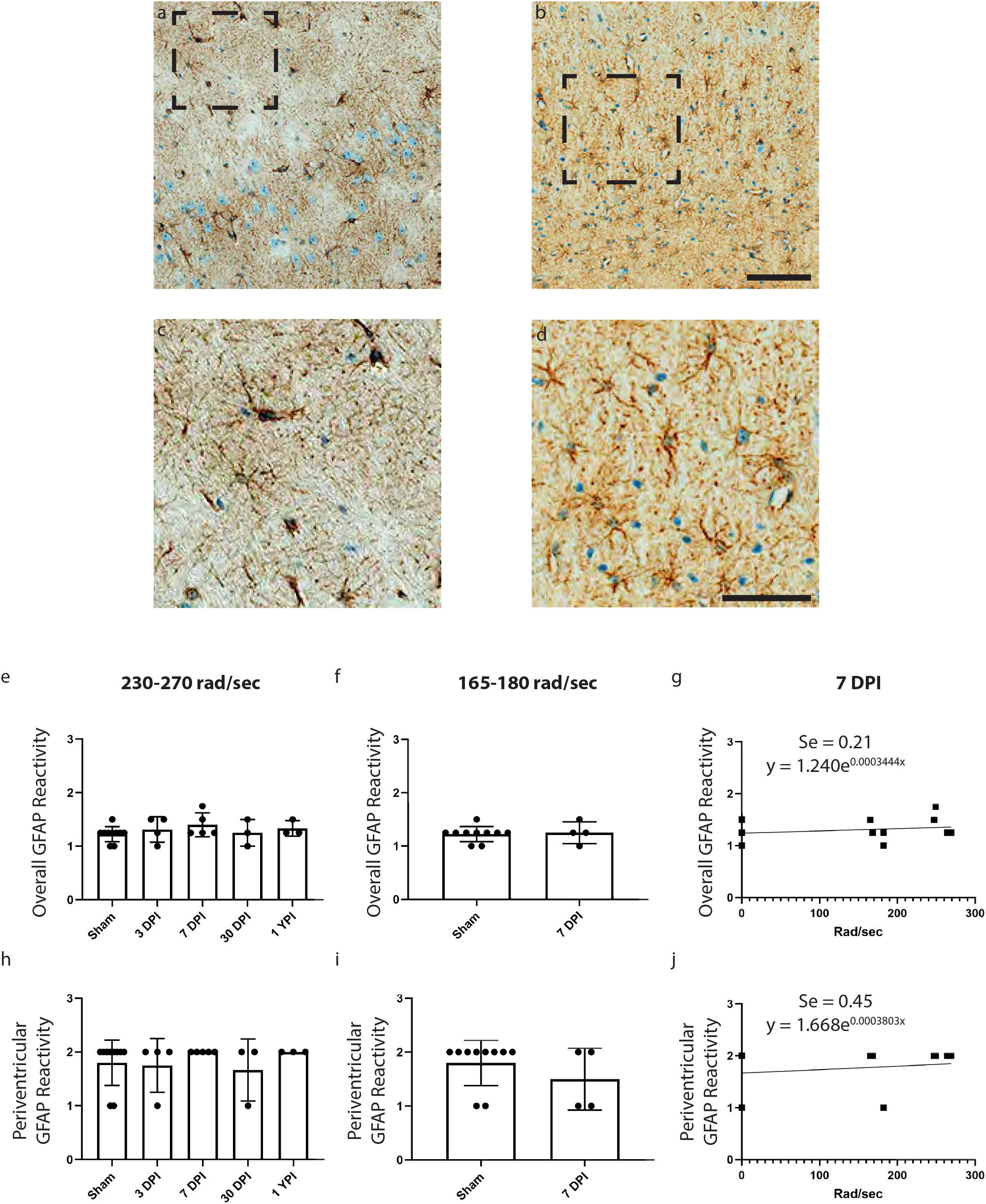
Examples of GFAP reactivity scored as a 1 (a) or 2 (b) are shown (scale = 100 μm) with corresponding call-out boxes for scores of 1(c) or 2(d; scale = 50 μm). No images were scored as a 3. There was no significant change in overall (e, f) or periventricular (h, i) GFAP reactivity in the 230-270 rad/sec or 165-185 rad/sec groups. Non-linear regression lines for 7 DPI specimens are shown (g, j).

For microglia cell density, Fiji software (National Institute of Health) was used to convert ionized calcium-binding adapter protein-1 (Iba-1) stained images to grayscale and perform color deconvolution, and then the “Analyze Particles” plugin was used to count cells in an automated fashion using an objective set of exclusion parameters (40). Particles less than 20 μm^2^ were excluded as these tended to be DAB stained microglial processes in the field of view, detached from a microglial cell body.

For Iba-1 skeletal analysis, we employed methods similar to Morrison et al. (33). Five 40x images per region of interest were examined using Fiji software. All Iba-1 positive cells in each 40x field were manually selected, and the image was deconvoluted. Bandpass filters, unsharp mask, and close plugins were applied before converting the image to binary, skeletonizing, and removing skeletons not overlaid with the manually selected cells (Supplementary Figure 1). The Analyze Skeleton plugin was then applied to quantify skeletal features such as number of process branches, junctions, process endpoints, and slab voxels in order to measure changes in microglia ramification (1). For each image, each feature was summed then divided by the total number of cells, thus providing a single field average normalized per cell. Slab voxels were then multiplied by the volume of the voxel to calculate the summed process length per cell.

For LFB analysis, measurements of the superior to inferior extent of the corpus callosum were obtained at 5 points along the mediolateral extent: at its juncture with the cingulate gyrus in both hemispheres as the lateral boundaries, at the midline of the corpus callosum, and midway between the corpus callosum midline and these lateral boundaries. These 5 measurements were averaged for each specimen. To measure the color intensity of the staining, the RGB color components were measured in ImageJ on a 0-255 AU scale. The scale for the blue color component was then inverted so that a zero value would indicate the whitest color while a 255 value would indicate the bluest color.

Statistical analysis was performed using GraphPad Prism statistical software (GraphPad Software Inc. La Jolla, CA). Due to low sample size, the 230-270 rad/sec injured group’s APP, GFAP, Iba-1 cell density, corpus callosum thickness, and LFB color intensity data were analyzed with a Kruskal-Wallis test and Dunn’s multiple comparisons. Kruskal-Wallis test results are reported as (*H* (degrees of freedom) = *H* test statistic, p-value). The 165-185 rad/sec injured group’s APP data were analyzed via a two-tailed Mann-Whitney test. Mann-Whitney results are reported as (*U* = U test statistic, p-value). Non-linear regression lines were created via an exponential growth equation. Goodness of fit is quantified using the standard deviation of the residuals (Se), the vertical distance (in Y units) of the experimental data point from the regression line, with a lower Se score indicative of a better predictive model. The skeletal analysis was statistically assessed via One-way analysis of variance (ANOVA) and Tukey’s multiple comparisons test. One-way ANOVA results are reported as (*F* (degrees of freedom numerator/degrees of freedom denominator) = *F* value, p-value). Mean, standard deviation, and 95% confidence intervals were reported. Differences were considered significant if p < 0.05. As this was an archival study, power calculations were not used to determine the number of specimens for each experimental group; the current study used all available specimens exposed to a single mild TBI. The number of images chosen for skeletal analysis was determined by power analysis from a pilot study.

## Results

### APP Pathology increased overall at 7 DPI but not at later time points

We performed semi-quantitative analysis of APP pathological burden in specific neuroanatomical areas in pig tissue sections at 3 DPI, 7 DPI, 30 DPI, and 1 YPI versus age-matched sham. There were no differences in any of the sham groups out to 1 YPI, so these animals were combined into a single sham group for statistical analysis. Previous literature has established APP pathology as the histological gold-standard to visualize axonal injury following closed-head TBI (for a review see (21)). Therefore, we sought to quantify the APP pathology burden in specimens subjected to closed-head rotational acceleration induced TBI at a range of rotational velocities that recreate the mechanistic forces and manifestations of mild TBI in humans. There was a significant increase in overall APP pathology burden in the 230-270 rad/sec injured group (*H* (4) = 11.75, p = 0.0193) at 7 DPI (Mean = 1.53, SD ± 0.30, 95% CI [1.16, 1.90]) compared to sham (Mean = 0.73, SD ± 0.42, 95% CI [0.43, 1.03]) (p = 0.0310) (Figure 1d). There were no significant changes to APP pathology at 3 DPI, 30 DPI, and 1 YPI compared to sham. Moreover, there was no change in the overall APP pathology burden in the 165-185 rad/sec injured group (*U* = 10, p = 0.1748) (Figure 1e). Pathology burden scoring was reliable between 2 trained observers (intraclass correlation coefficient = 0.834). Examining all 7 DPI specimens and age-matched shams across all injury levels, a non-linear regression line was found (Se = 0.28), depicting the non-linear increase in global APP pathology as injury level increased (Figure 1f).

### APP Pathology increased in the periventricular white matter at 3 DPI and 7 DPI, and in the fimbria/fornix at 7 DPI, but not at later time points

Additionally, we analyzed the APP pathology burden in each examined neuroanatomical region to detect areas potentially more vulnerable to axonal pathology in our model. In the periventricular white matter, there was a significant increase in APP pathology burden in the 230-270 rad/sec injured group (*H* (4) = 18.82, p = 0.0009) at 7 DPI (Mean = 2.2, SD ± 0.45, 95% CI [1.65, 2.76]) compared to sham (Mean = 0.40, SD ± 0.52, 95% CI [0.03, 0.77]) (p = 0.0008), as well as 3 DPI (Mean = 1.75, SD ± 0.50, 95% CI [0.95, 2.55]) compared to sham (p = 0.0372) (Figure 1g). Like our overall APP pathology data, there were no significant changes at 30 DPI and 1 YPI. There was no change in the APP pathology burden in the 165-185 rad/sec injured group (*U* = 8, p = 0.0849) (Figure 1h). Examining all 7 DPI specimens and age-matched sham across all injury levels, a non-linear regression line was found (Se = 0.43), illustrating the exponential increase in periventricular white matter APP pathology burden according to injury level (Figure 1i).

While subsequent analyses focus on the hippocampus, we also examined the fimbria/fornix region surrounding the hippocampus for APP pathology. In the fimbria/fornix, there was a significant increase in APP pathology burden in the 230-270 rad/sec injured group (*H* (4) = 13.00, p = 0.0113) at 7 DPI (Mean = 1.2, SD ± 0.45, 95% CI [0.64, 1.76]) compared to sham (Mean = 0.10, SD ± 0.32, 95% CI [−0.13, 0.33]) (p = 0.0072) (Supplementary Figure 2a). There were no significant changes to APP pathology at 30 DPI and 1 YPI compared to sham. There was no change in the APP pathology burden in the 165-185 rad/sec injured group (*U* = 17, p = 0.9999) (Supplementary Figure 2b). Examining all 7 DPI specimens and age-matched sham across all injury levels, a non-linear regression line was found (Se = 0.36), displaying the exponential increase in fimbria/fornix APP pathology burden according to injury level (Supplementary Figure 2c).

The striatum, dorsal and ventral thalamus, and cerebellum were also assessed for axonal pathology at all time points and injury levels. However, no significant changes in APP pathology burden were observed in these areas (Supplementary Figure 2d-o).

### GFAP reactivity did not change after single mild TBI

Reactive astrogliosis is often used as a marker for damaged CNS tissue and exists on a continuum of genetic and cellular changes (44). There was no significant change in overall GFAP reactivity in the 230-270 rad/sec injured group (*H* (4) = 2.846, p = 0.5839) (Figure 2c), nor was there a change in the overall GFAP reactivity in the 165-185 rad/sec injured group (*U* = 18.50, p = 0.9970) (Figure 2d).

Moreover, there was no significant change in periventricular white matter GFAP reactivity in the 230-270 rad/sec injured group (*H* (4) = 2.452, p = 0.6532) (Figure 2f), nor was there a change in the overall GFAP reactivity in the 165-185 rad/sec injured group (*U* = 14, p = 0.5205) (Figure 2g).

### Microglia skeletal analysis revealed morphological changes in the periventricular white matter at 7 DPI

Microglia have been shown to alter their morphology in response to a change in CNS homeostasis (38). Therefore, we quantified potential morphological changes in anatomical areas with axonal pathology via automated skeletal analyses (Supplementary Figure 1). In the periventricular white matter, there were significant increases in the number of process branches, junctions, and endpoints per microglia. Specifically, there was an increase in the number of branches per cell (*F* (4/120) = 3.402, p = 0.0113) at 7 DPI (Mean = 21.81, SD ± 7.23, 95% CI [18.83, 24.80]) compared to sham (Mean = 14.9, SD ± 7.24, 95% CI [12.85, 16.96]) (p = 0.0184) (Figure 3h). There was also a significant change in the number of junctions per microglia (*F* (4/120) = 3.310, p = 0.0131) with an increase at 7 DPI (Mean = 10.46, SD ± 3.65, 95% CI [8.95, 11.96]) compared to sham (Mean = 7.00, SD ± 3.70, 95% CI [5.95, 8.05]) (p = 0.0223) (Figure 3i). Additionally, there was an increase in the number of endpoints per microglia (*F* (4/120) = 3.588, p = 0.0084) with an increase at 7 DPI (Mean = 11.33, SD ± 3.20, 95% CI [10.01, 12.65]) compared to sham (Mean = 8.31, SD ± 3.11, 95% CI [7.42, 9.19]) (p = 0.0117) (Figure 3j). However, there were no significant changes in branches, junctions, or endpoints at 30 DPI or 1 YPI, and there was no significant change in summed process length per microglia (*F* (4/120) = 2.355, p = 0.0576) (Figure 3k).

**Figure 3.**
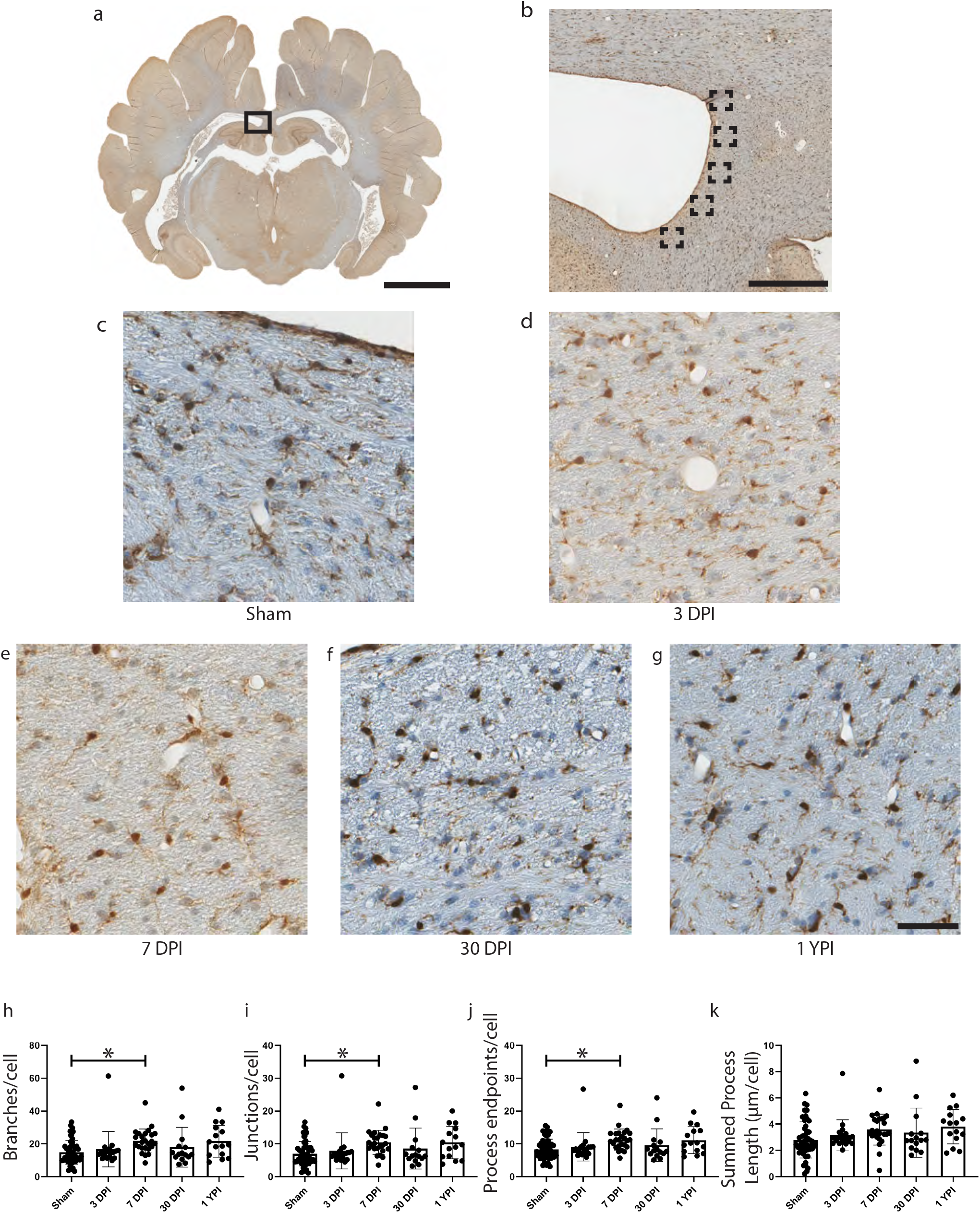
A whole coronal section of Iba-1 stained tissue is shown with a call-out box centered around the periventricular white matter (a, scale = 1 cm). Five call-out boxes in the periventricular white matter depict the location of the 40x images used for skeletal analysis (b, scale = 500 μm). 40x images for all injury levels are displayed (c-g, scale = 50 μm). There is a significant increase in the number of branches (p = 0.0184), junctions (p = 0.0131), and process endpoints (p = 0.0223) per cell at 7 DPI compared to sham.

### Microglia skeletal analysis revealed subacute changes in the anterior hippocampal hilus

In anterior hippocampal hilus, we observed significant increases to the number of branches, junctions, and endpoints per cell at 3 DPI and 7 DPI, and an increase in summed process length at 3 DPI (Table 1). Specifically, there was a significant increase in the number of branches per cell (*F* (4/119) = 4.171, p = 0.0034) at 7 DPI (p = 0.0260) and 3 DPI (p = 0.0124) compared to sham (Figure 4d). The number of junctions per cell significantly increased (*F* (4/119) = 4.104, p = 0.0038) at 7 DPI (p = 0.0324) and 3 DPI (p = 0.0116) compared to sham (Figure 4e). Additionally, the number of endpoints per microglia significantly increased (*F* (4/119) = 4.389, p = 0.0024) at 7 DPI (p = 0.0165) and 3 DPI (p = 0.0128) compared to sham (Figure 4f). Finally, the summed process length per cell increased (*F* (4/119) = 3.854, p = 0.0056) at 3 DPI compared to sham (p = 0.0076) (Figure 4g). There were no significant changes at 30 DPI or 1 YPI.

**Table 1.**
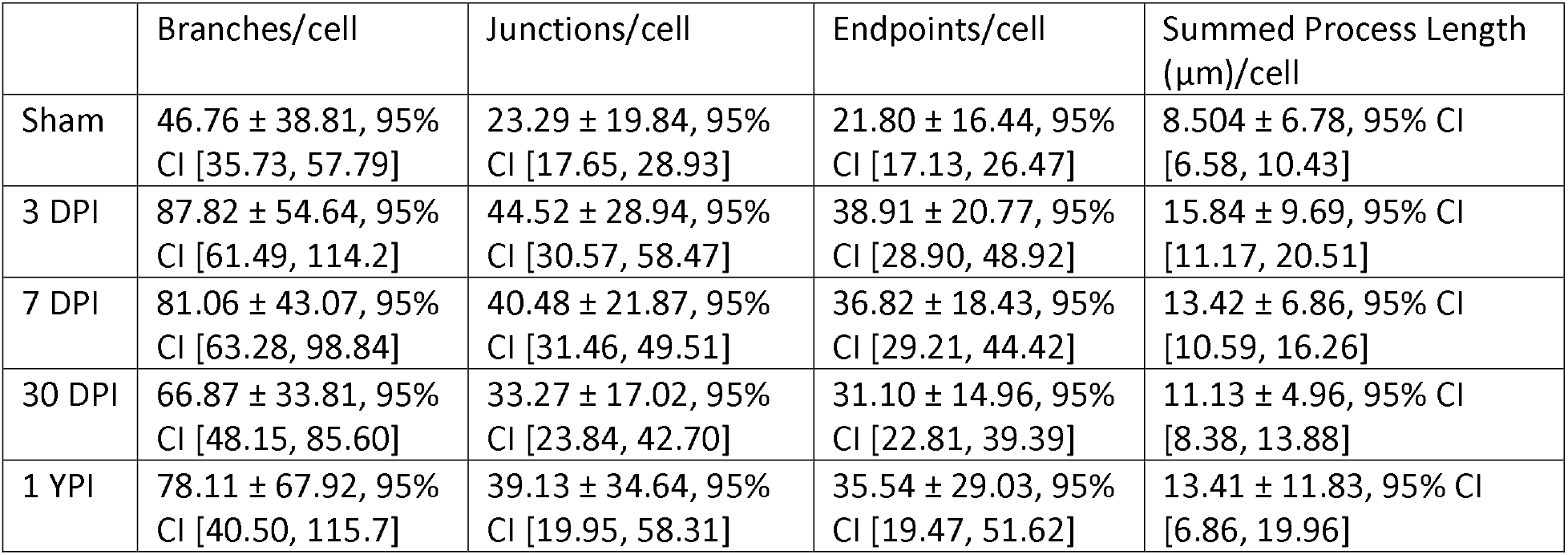
Skeletal analysis features in anterior hippocampal hilus. All values are reported as mean ± standard deviation, 95% confidence interval [Lower 95% CI, Upper 95% CI].

**Figure 4.**
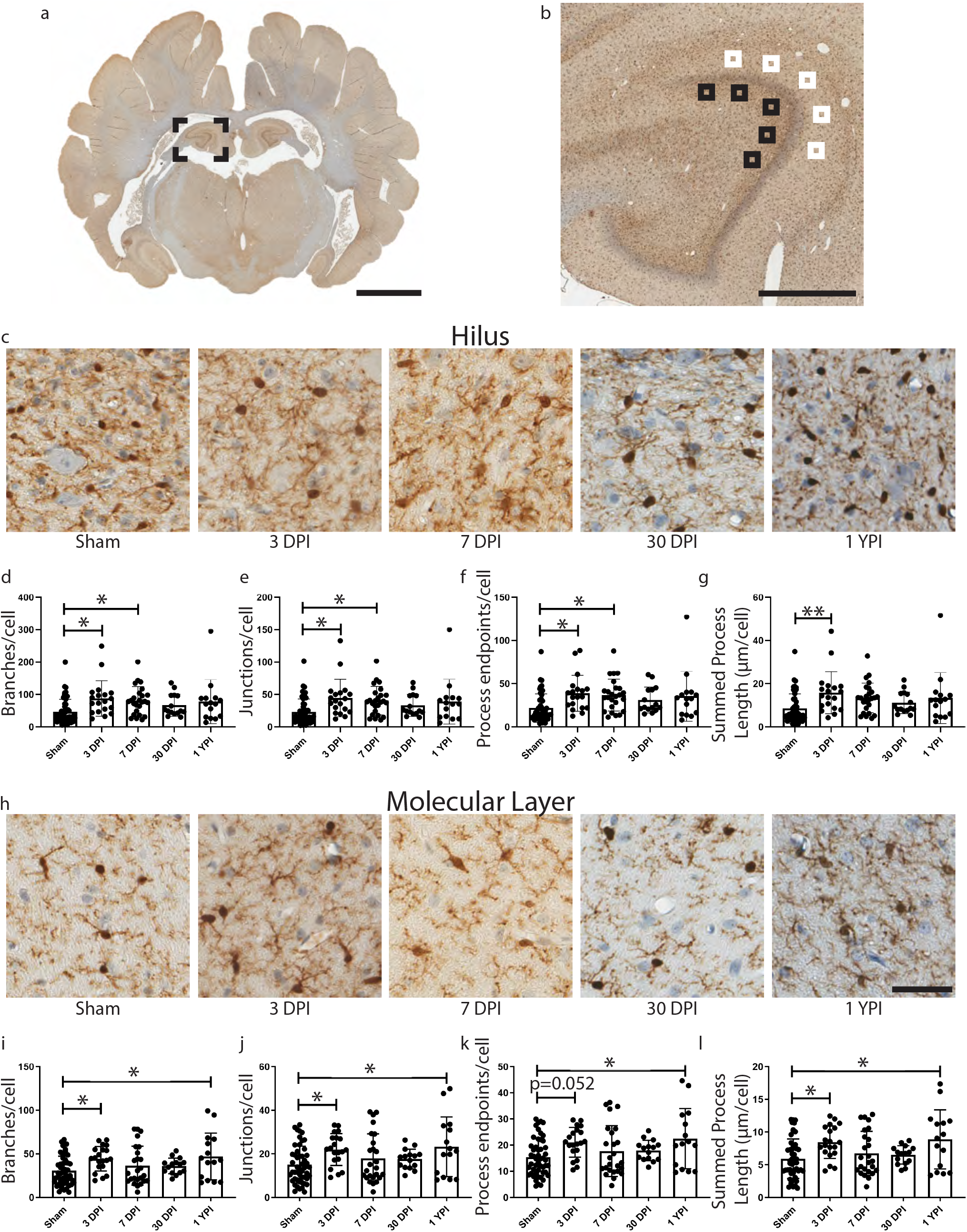
A whole coronal section of Iba-1 stained tissue is shown with a call-out box centered around the dorsal aspect of anterior hippocampus (a, scale = 1 cm). Five white call-out boxes in the molecular layer and five black call-out boxes in the hilus depict the location of the 40x images used for skeletal analysis (b, scale = 1 mm). 40x images in the hilus for all injury levels are displayed (c). There is a significant increase in the number of branches (p = 0.0260), junctions (p = 0.0324), and process endpoints (p = 0.0165) per cell in the hilus at 7 DPI compared to sham. Additionally, there is a significant increase in the number of branches (p = 0.0124), junctions (p = 0.0116), process endpoints (p = 0.0128), and summed process length (p = 0.0076) per cell in the hilus at 3 DPI compared to sham. 40x images in the molecular layer for all injury levels are displayed (h; scale = 50 μm). There is a significant increase in the number of branches (p = 0.0462), junctions (p = 0.0450), and summed process length (p = 0.0231) per cell in the molecular layer at 3 DPI compared to sham. Additionally, there is a significant increase in the number of branches (p = 0.0260), junctions (p = 0.0286), process endpoints (p = 0.0217), and summed process length (p = 0.0157) per cell in the molecular layer at 1 YPI compared to sham.

### Microglia skeletal analysis revealed chronic changes in the anterior hippocampal molecular layer

In anterior hippocampal molecular layer, we observed significant increases to the number of branches, junctions, and the summed process length per cell at 3 DPI, and an increase in the branches, junctions, endpoints, and summed process length at 1 YPI (Table 2). There was a significant increase in the number of branches per cell (*F* (4/120) = 3.371, p = 0.0119) at 3 DPI (p = 0.0462) and 1 YPI (p = 0.0260) compared to sham (Figure 4i). The number of junctions per cell significantly increased (*F* (4/120) = 3.344, p = 0.0038) at 3 DPI (p = 0.0450) and 1 YPI (p = 0.0450) compared to sham (Figure 4j). Furthermore, the number of endpoints per microglia significantly increased (*F* (4/120) = 3.396, p = 0.0114) at 1 YPI (p = 0.0217) compared to sham (p = 0.0217), yet did not change at 3 DPI compared to sham (p = 0.0523) (Figure 4k). Finally, the summed process length per cell significantly increased (*F* (4/120) = 4.082, p = 0.0039) at 3 DPI (p = 0.0231) and 1 YPI (p = 0.0157) compared to sham (Figure 4l).

**Table 2.** Skeletal analysis features in anterior hippocampal molecular layer. All values are reported as mean ± standard deviation, 95% confidence interval [Lower 95% CI, Upper 95% CI].

| | Branches/cell | Junctions/cell | Endpoints/cell | Summed Process Length ($\mu\text{m}$)/cell |
| --- | --- | --- | --- | --- |
| Sham | 30.78 ± 16.58, 95% CI [26.07, 35.49] | 15.04 ± 8.49, 95% CI [12.63, 17.46] | 15.35 ± 7.10, 95% CI [13.34, 17.37] | 5.92 ± 3.03, 95% CI [5.06, 6.78] |
| 3 DPI | 44.37 ± 14.07, 95% CI [37.79, 50.96] | 22.00 ± 7.30, 95% CI [18.58, 25.41] | 21.08 ± 5.69, 95% CI [18.41, 23.74] | 8.44 ± 2.48, 95% CI [7.28, 9.60] |
| 7 DPI | 36.46 ± 22.31, 95% CI [27.25, 45.67] | 17.92 ± 11.27, 95% CI [13.27, 22.57] | 17.72 ± 9.71, 95% CI [13.71, 21.73] | 6.74 ± 3.37, 95% CI [5.35, 8.13] |
| 30 DPI | 36.12 ± 8.95, 95% CI [31.16, 41.07] | 17.60 ± 4.52, 95% CI [15.10, 20.11] | 17.99 ± 3.88, 95% CI [15.83, 20.14] | 6.50 ± 1.40, 95% CI [5.72, 7.28] |
| 1 YPI | 47.04 ± 26.89, 95% CI [32.15, 61.93] | 23.24 ± 13.68, 95% CI [15.66, 30.81] | 22.46 ± 11.50, 95% CI [16.09, 28.83] | 8.87 ± 4.52, 95% CI [6.37, 11.37] |

### Microglia skeletal analysis of posterior hippocampus revealed subacute changes in the hilus and molecular layer

In posterior hippocampal hilus, we observed significant increases to the number of branches, junctions, endpoints, and summed process length per cell at 7 DPI and 30 DPI (Table 3). There was a significant increase in the number of branches per cell (*F* (4/120) = 6.483, p < 0.001) at 7 DPI (p = 0.0011) and 30 DPI (p = 0.0085) compared to sham (Figure 5d). The number of junctions per cell significantly increased (*F* (4/120) = 6.446, p < 0.0001) at 7 DPI (p = 0.0010) and 30 DPI (p = 0.0094) compared to sham (Figure 5e). Additionally, the number of endpoints per microglia significantly increased (*F* (4/120) = 6.383, p = 0.0001) at 7 DPI (p = 0.0016) and 30 DPI (p = 0.0069) compared to sham (Figure 5f).The summed process length per cell increased (*F* (4/120) = 5.028, p = 0.0009) at 7 DPI (p = 0.0095) and 30 DPI (p = 0.0223) compared to sham (Figure 5g).

**Table 3.** Skeletal analysis features in posterior hippocampal hilus. All values are reported as mean ± standard deviation, 95% confidence interval [Lower 95% CI, Upper 95% CI].

| | Branches/cell | Junctions/cell | Endpoints/cell | Summed Process Length ($\mu\text{m}$)/cell |
| --- | --- | --- | --- | --- |
| Sham | 53.35 ± 27.91, 95% CI [45.42, 61.29] | 26.58 ± 14.25, 95% CI [22.53, 30.63] | 24.81 ± 12.02, 95% CI [21.39, 28.22] | 9.75 ± 4.90, 95% CI [8.36, 11.14] |
| 3 DPI | 50.46 ± 26.47, 95% CI [38.07, 62.84] | 25.16 ± 13.66, 95% CI [18.77, 31.56] | 23.45 ± 11.02, 95% CI [18.29, 28.61] | 9.18 ± 4.95, 95% CI [6.87, 11.50] |
| 7 DPI | 88.09 ± 54.43, 95% CI [65.62, 110.6] | 44.29 ± 27.58, 95% CI [32.91, 55.68] | 39.28 ± 23.38, 95% CI [29.62, 48.93] | 14.59 ± 8.59, 95% CI [11.05, 18.14] |
| 30 DPI | 88.29 ± 38.77, 95% CI [66.83, 109.8] | 44.19 ± 19.58, 95% CI [33.34, 55.03] | 40.09 ± 16.92, 95% CI [30.72, 49.46] | 14.99 ± 5.74, 95% CI [11.81, 18.17] |
| 1 YPI | 65.91 ± 21.61, 95% CI [53.94, 77.88] | 33.17 ± 11.10, 95% CI [27.02, 39.31] | 29.57 ± 9.04, 95% CI [24.57, 34.57] | 11.90 ± 3.92, 95% CI [9.73, 14.07] |

**Figure 5.**
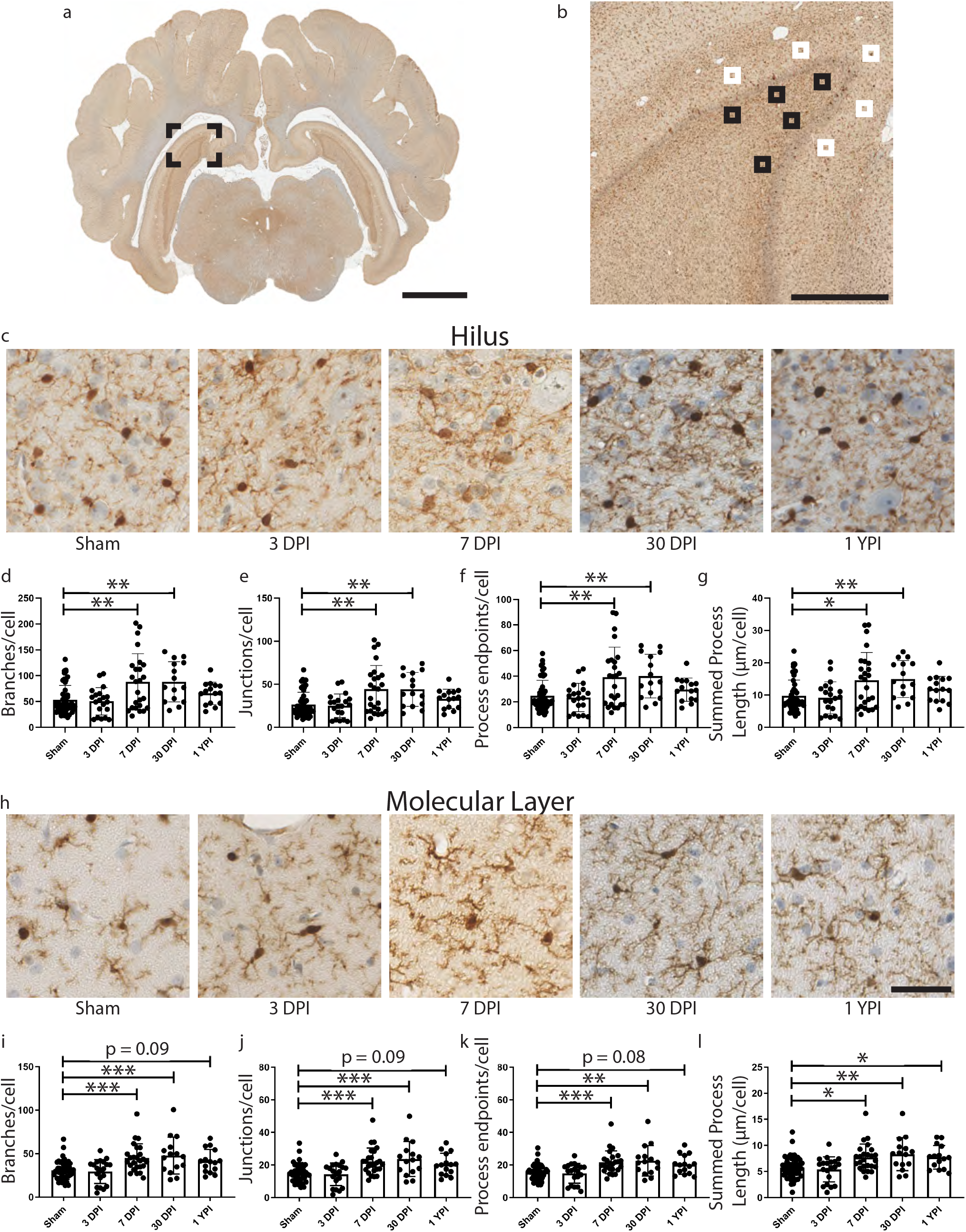
A whole coronal section of Iba-1 stained tissue is shown with a call-out box centered around the dorsal aspect of posterior hippocampus (a, scale = 1 cm). Five white call-out boxes in the molecular layer and five black call-out boxes in the hilus depict the location of the 40x images used for skeletal analysis (b, scale = 1 mm). 40x images in the hilus for all injury levels are displayed (c). There is a significant increase in the number of branches (p = 0.0011), junctions (p = 0.0010), process endpoints (p = 0.0016), and summed process length (p = 0.0095) per cell in the hilus at 7 DPI compared to sham. Additionally, there is a significant increase in the number of branches (p = 0.0085), junctions (p = 0.0094), process endpoints (p = 0.0069), and summed process length (p = 0.0223) per cell in the hilus at 30 DPI compared to sham. 40x images in the molecular layer for all injury levels are displayed (h; scale = 50 μm). There is a significant increase in the number of branches (p = 0.0007), junctions (p = 0.0008), process endpoints (p = 0.0009), and summed process length (p = 0.0126) per cell in the molecular layer at 7 DPI compared to sham. Additionally, there is a significant increase in the number of branches (p = 0.0008), junctions (p = 0.0007), process endpoints (p = 0.0016), and summed process length (p = 0.0044) per cell in the molecular layer at 30 DPI compared to sham. Finally, there is a significant increase in summed process length at 1 YPI compared to sham (p =0.0316).

In posterior hippocampal molecular layer, we observed significant increases to the number of branches, junctions, endpoints, and summed process length per cell at 7 DPI and 30 DPI (Table 4). Moreover, we found a significant increase in summed process length at 1 YPI. Specifically, there was a significant increase in the number of branches per cell (*F* (4/120) = 8.361, p < 0.001) at 7 DPI (p = 0.0007) and 30 DPI (p = 0.0008)compared to sham (Figure 5i).The number of junctions per cell significantly increased (*F* (4/120) = 8.347, p < 0.0001) at 7 DPI (p = 0.0008) and 30 DPI (p = 0.0007) compared to sham (Figure 5j). Additionally, the number of endpoints per microglia significantly increased (*F* (4/120) = 7.954, p > 0.0001) at 7 DPI (p = 0.0009) and 30 DPI (p = 0.0016) compared to sham (Figure 5k).The summed process length per cell increased (*F* (4/120) = 6.736, p < 0.0001) at 7 DPI (p = 0.0126), 30 DPI (p = 0.0044), and 1 YPI (p = 0.0316) compared to sham (Figure 5l). This increase in summed process length is the only statistically significant change at 1 YPI in the posterior hippocampus.

**Table 4.** Skeletal analysis features in posterior hippocampal molecular layer. All values are reported as mean ± standard deviation, 95% confidence interval [Lower 95% CI, Upper 95% CI].

| | Branches/cell | Junctions/cell | Endpoints/cell | Summed Process Length ($\mu\text{m}$)/cell |
| --- | --- | --- | --- | --- |
| Sham | 30.76 ± 10.53, 95% CI [27.77, 33.75] | 14.99 ± 5.38, 95% CI [13.46, 16.52] | 15.50 ± 4.50, 95% CI [14.22, 16.78] | 5.78 ± 2.05, 95% CI [5.19, 6.36] |
| 3 DPI | 29.45 ± 13.75, 95% CI [23.02, 35.88] | 14.28 ± 6.94, 95% CI [11.04, 17.53] | 14.90 ± 6.08, 95% CI [12.06, 17.75] | 5.42 ± 2.40, 95% CI [4.30, 6.54] |
| 7 DPI | 45.03 ± 16.22, 95% CI [38.34, 51.73] | 22.15 ± 8.26, 95% CI [18.74, 25.56] | 21.62 ± 7.01, 95% CI [18.73, 24.52] | 7.67 ± 2.60, 95% CI [6.60, 8.75] |
| 30 DPI | 47.82 ± 21.26, 95% CI [36.05, 59.59] | 23.70 ± 10.71, 95% CI [17.76, 29.63] | 22.56 ± 9.34, 95% CI [17.39, 27.73] | 8.33 ± 3.16, 95% CI [6.58, 10.07] |
| 1 YPI | 41.43 ± 13.43, 95% CI [33.99, 48.87] | 20.37 ± 6.78, 95% CI [16.61, 24.12] | 20.17 ± 5.90, 95% CI [16.90, 23.44] | 7.74 ± 2.26, 95% CI [6.49, 8.99] |

### Microglia skeletal analysis features changed after 3 DPI, but microglia cell density does not change after single mild TBI in the corpus callosum

In the corpus callosum, there were no significant changes in skeletal analysis features at time points post injury compared to sham. However, there were several significant changes compared to 3 DPI time points.

There was a significant increase in the number of branches per cell (*F* (4/120) = 4.685, p = 0.0015) at 7 DPI (Mean = 26.34, SD ± 8.95, 95% CI [22.64, 30.03]) compared to 3 DPI (Mean = 18.85, SD ± 3.14, 95% CI [17.38, 20.32] (p = 0.0049), as well as a significant increase at 1 YPI (Mean = 27.59, SD ± 8.00, 95% CI [23.15, 32.02]) compared to 3 DPI (p = 0.0040) (Figure 6i). There was a significant increase in the number of junctions per cell (*F* (4/120) = 4.624, p = 0.0017) at 7 DPI (Mean = 12.89, SD ± 4.61, 95% CI [10.99, 14.79]) compared to 3 DPI (Mean = 9.02, SD ± 1.62, 95% CI [8.26, 9.78] (p = 0.0050), as well as a significant increase at 1 YPI (Mean = 13.55, SD ± 4.12, 95% CI [11.26, 15.83]) compared to 3 DPI (p = 0.0040) (Figure 6j). Additionally, there was a significant increase in the number of endpoints per cell (*F* (4/120) = 4.757, p = 0.0013) at 7 DPI (Mean = 13.13, SD ± 3.72, 95% CI [11.60, 14.67]) compared to 3 DPI (Mean = 10.09, SD ± 1.33, 95% CI [9.47, 10.71] (p = 0.0053), as well as a significant increase at 1 YPI (Mean = 13.62, SD ± 3.26, 95% CI [11.82, 15.42]) compared to 3 DPI (p = 0.0047) (Figure 6k). It should be noted that the branches, junctions, and endpoints analyses did not pass a test for equal variance according to Brown-Forsythe tests, and therefore these populations have significantly different standard deviations. Finally, there was a significant change overall in the summed process length per cell (*F* (4/120) = 2.537, p = 0.0435) but there was no statistical difference between any two experimental groups by post hoc analysis (Figure 6l).

**Figure 6.**
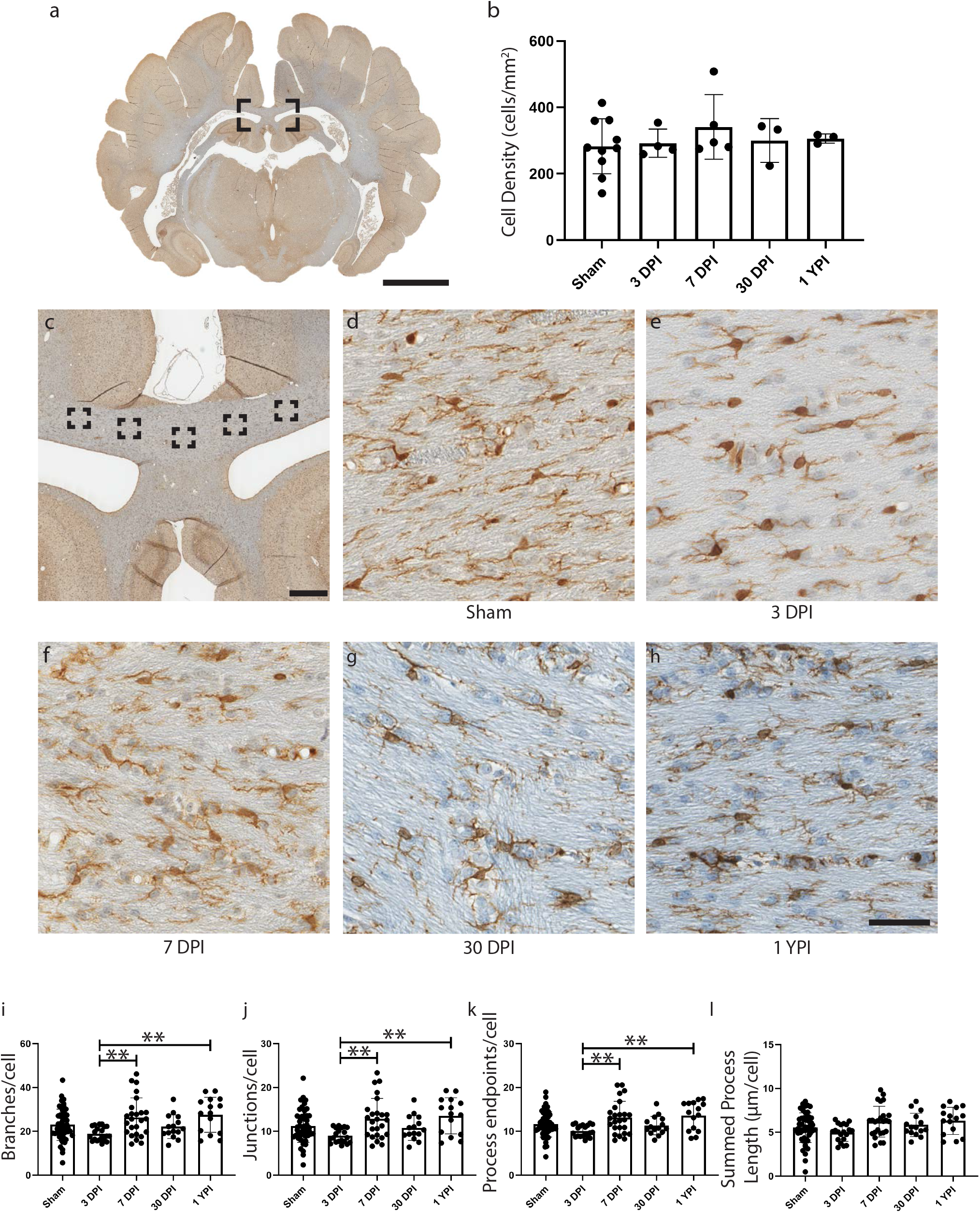
A whole coronal section of Iba-1 stained tissue is shown with a call-out box centered around the corpus callosum (a, scale = 1 cm). The density of Iba-1+ cells did not change in the corpus callosum at time points post-injury (b). Five call-out boxes in the corpus callosum depict the location of the 40x images used for skeletal analysis (c, scale = 1 mm). 40x images for all injury levels are displayed (d-h, scale = 50 μm). There is no significant change in injured specimen’s skeletal analysis features compared to sham. However, there is a significant increase in the number of branches (p = 0.0049), junctions (p = 0.0050), and process endpoints (p = 0.0053) per cell at 7 DPI compared to 3 DPI. There is also a significant increase in the number of branches (p = 0.0040), junctions (p = 0.0040), and process endpoints (p = 0.0047) per cell at 1 YPI compared to 3 DPI.

In a previous study from our group, we conducted microglia cell density analysis in discrete subregions of the hippocampus (17). Here, we applied that same analysis to the corpus callosum; however, we did not detect any changes in microglia density following injury at any time point (*H* (4) = 1.520, p = 0.8231).

### Corpus callosum thickness and LFB staining intensity did not change following single mild TBI

Histopathological studies in humans have noted white matter degeneration and corpus callosum thinning at greater than 1 YPI after a single moderate to severe TBI (20). Therefore, we sought to determine if corpus callosum thickness changes after a single mild TBI in our pig model of injury. We did not find a significant change in corpus callosum thickness at any time point after a single mild TBI (*H* (4) = 5.070, p = 0.2802) (Figure 7g). Additionally, we examined if the intensity of LFB staining changes after injury, potentially indicating myelin loss. Again, no significant change was detected (*H* (4) = 8.715, p = 0.0686) (Figure 7h).

**Figure 7.**
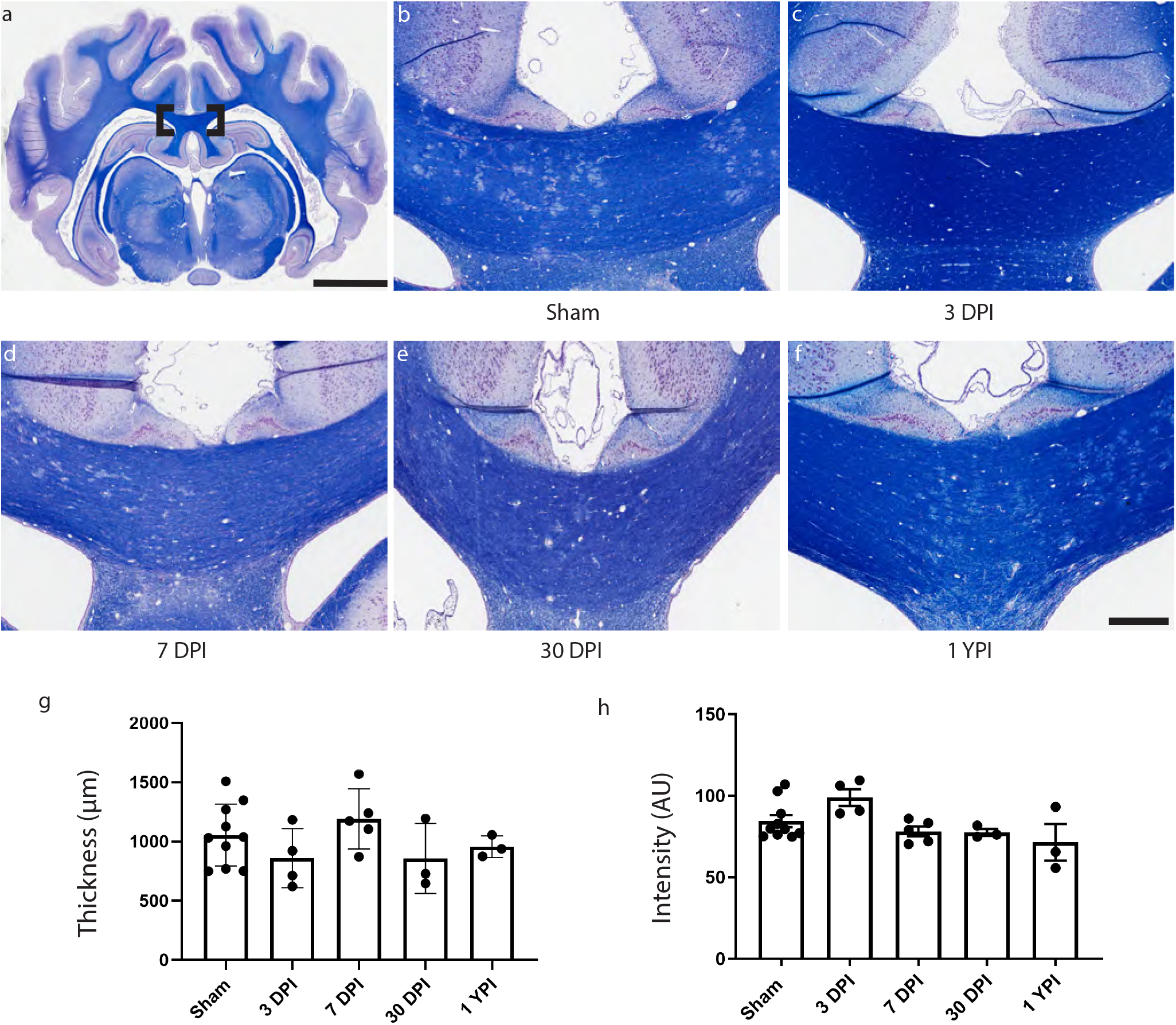
A whole coronal section of LFB stained tissue is shown with a call-out box centered around the corpus callosum (a, scale = 1 cm). Representative LFB staining is shown in the corpus callosum for all experimental groups (b-f; scale = 1 mm). Neither the thickness of the corpus callosum (g) or the staining intensity of the LFB (h) significantly changed as a result of injury.

## Discussion

A history of mild TBI is associated with chronic health problems such as reduced memory performance and an increased likelihood of neurological disorders, as well as life-long healthcare costs (12,19,32). After a single mild TBI caused by head rotation at 230-270 rad/sec using our pig model of closed-head diffuse brain injury, we found significant increases in APP pathology burden at 3 DPI and 7 DPI, which resolved by 30 DPI. Importantly however, we did find microglial morphological changes out to chronic time points, indicating a more ramified phenotype and possible altered function. Specifically, we found an increase in the number of branches, junctions, endpoints, and summed process length per microglia at acute and subacute time points in the hippocampal hilus, and at acute, subacute, and chronic time points in the hippocampal molecular layer. Moreover, we found an increase in the number of branches, junctions, and endpoints in the periventricular white matter at 7 DPI, which coincides with an increase in APP pathology. These results support our original hypothesis that acute axonal pathology and regional microglial morphological changes will occur after a single mild TBI. These morphological changes coincide with our previous study that demonstrated microglia density increases in hippocampal subregions after a single mild TBI (17).

In the current study, we first assessed the APP pathology burden within the tissue, as APP accumulations are the hallmark pathology of TBI and axonal injury. Following axonal injury, anterograde vesicular transport along the axon is disrupted and APP accumulates intracellularly (14,21). Traditionally, this protein buildup will progress from hours to months post injury, the distal axonal segment will gradually undergo Wallerian degeneration, and the cellular debris will be phagocytosed primarily by microglia (2,25). These APP+ axons can therefore take the form of discrete bulb formations at the end of disconnected axons or varicosities along the length of an axon depending on its time course through this degenerative process. APP pathology has historically been characterized in this pig model of injury (3). However, we used a new strain of pig that has a lower peak body weight and is more amenable to animal handling out to 1 year following injury. Therefore, re-characterization of APP pathology was necessary. We observed an increase in the overall APP pathology burden at 7 DPI at forces of 230-270 rad/sec and no change at forces of 165-185 rad/sec (Figure 1). This may inform future rotational injury studies of the forces needed to obtain significant axonal injury. Moreover, when we assessed specific neuroanatomical regions for APP pathology, we noted an increase at 3 DPI and 7 DPI in the periventricular white matter and at 7 DPI in the fimbria/fornix surrounding the hippocampus. This is consistent with previous pathological assessments of this injury model. Browne et al. created regional schematics of APP+ axons at 7 DPI in both single coronal and axial rotational injuries, illustrating axonal pathology within the deep white matter (3). Importantly, we examined APP pathology out to 1 YPI, but did not observe significant pathology at this chronic time point, thus suggesting significant clearance of damaged axons by phagocytosis in our model. Yet this does not necessarily mean that all damaged axons are phagocytosed. Some human post-mortem evidence suggests that axonal degeneration may occur for years after injury, though the degree of axonal pathology is less extensive than acute axonal pathology (6). Additionally, APP staining for axonal injury may not identify all injured axons. Subpopulations of injured axons may undergo calcium-dependent proteolysis of the cytoskeleton, while other axons may experience physiological changes such as increased intra-axonal calcium or sodium (23,56,57). Neuronal plasma membrane disruptions may also occur causing a loss of ionic gradients and osmotic imbalance (8,54). Future studies utilizing multiple markers for trauma-induced alterations will identify a more comprehensive profile of mild TBI pathology, potentially informing clinical outcomes.

We also examined the changes to microglial morphology in anatomical subregions in response to mild TBI. We began with the periventricular white matter, where we found an increase in the number of microglial branches, junctions, and endpoints at 7 DPI. While microglia morphological changes have been noted following injury in many studies after TBI, microglia skeletal analysis has become increasingly utilized for in-depth quantitative morphological analysis, particularly when subtle changes are suspected after mild injuries (33,45,46). Morrison et al. employed skeletal analysis in a rodent model of fluid percussion injury, finding a decrease in microglia ramification as measured by summed process length and number of endpoints around the site of impact (33). In contrast we found an increase in microglia ramification in a region vulnerable to TBI. These conflicting responses may be due to the different species and injury model used in the current study versus the Morrison study. Gorse and Lafrenaye explored these species differences in neuro-immune interactions while characterizing the interaction of APP+ axons with microglia in both rats and pigs at 6 hours post injury and 1 DPI; rats had a reduction in microglia-to-injured axon interactions at 6 hours while pigs had an increase at 1 DPI (16). While the current study did not examine the direct interactions of microglia and APP+ axons, we did detect an increase in microglia ramification in a region with significantly increased APP pathology. Future studies in our model could examine the role of microglia around APP pathology to determine if microglia are adopting a more phagocytic behavior, potentially contributing to synapse remodeling, or promoting neurotoxicity by increasing inflammatory cytokine release (38,52).

Microglia morphological changes were also seen in subregions of the hippocampus. It is important to note that we did not assess APP pathology in the hippocampus, but rather in the fimbria/fornix, the main efferent white matter tract from the hippocampus, which also contains some afferent projections to the hippocampus. The hippocampus, a structure essential for memory formation, has been reported to have synaptic dysfunction and neuropathological alterations after a single mild TBI in both rat and pig preclinical models (17,30,39,43,55). In a previous study, we detailed an increase in microglia density in hippocampal subregions after injury, particularly in the molecular layer (17). We expanded on that microglial analysis in the current study by using skeletal analysis. In both anterior and posterior hippocampal hilus, we found various morphological changes out to 30 DPI. Yet, in both anterior and posterior hippocampal molecular layer, we found morphological changes out to 1 YPI. This suggests that the molecular layer may be vulnerable to chronic inflammation after a single mild TBI.

Unfortunately, we are limited in the interpretation of these morphological changes at this time. Traditional microglial evaluation attempted to classify resting or inactivated microglia as ramified whereas activated microglia are ameboid. While phagocytic microglia tend to exhibit an ameboid morphology, ramified microglia are also active and continually survey the parenchyma by clearing metabolic products and tissue debris, as well as monitoring the functional status of synapses (34,51). Additionally, transcriptomic profiling of microglia has shown a much more dynamic response to injury: microglial changes are context dependent and cannot fit into a classic M1 (pro-inflammatory) or M2 (anti-inflammatory) bimodal arrangement (35,53). Therefore, morphological changes in microglia likely indicate that microglia have detected a change in homeostasis (38), and it appears that microglia in our model register a homeostatic change in the molecular layer out to 1 YPI. It is possible that our observed microglial changes indicate increased surveillance for debris and synapse functionality, but additional experiments to assess neuronal and synaptic integrity are needed to understand this complicated response.

Moreover, our molecular layer skeletal analysis examined photomicrographs immediately superior to the granule cell layer. This region is where the commissural/associational fibers terminate from the hilar mossy cells. While we did not find chronic hilar changes, efferent fibers from hilar mossy cells may still be chronically altered in the molecular layer. Future experiments should consider both whole cell electrophysiological recordings on mossy cells, as well as local field potential recordings in the molecular layer, at chronic time points to better characterize functional changes to the hippocampus. Circuit level disruption of the hippocampus has been reported from our group in ex vivo hippocampal slices and presynaptic staining around mossy cells have increased at 7 DPI (17,55). The synthesis of this physiological and histological data suggests that the next steps should be an examination of the mossy cell outputs.

We also assessed GFAP reactivity following injury but did not observe any significant changes at any time point post-injury compared to sham. This is particularly interesting as reactive astrocytes are widely considered a hallmark pathology of diseased or injured CNS tissue (44). In the healthy brain, astrocytes maintain the extracellular space by buffering neurotransmitters and regulating osmolarity; however in TBI pathogenesis, astrocytes can undergo genomic and proliferative changes that may vary depending on the severity of the injury and location in the brain (for a review, see (4)). Within this model of closed-head, single mild TBI, colleagues have described blood-brain barrier breakdown through the histological colocalization of fibrinogen and astrocytes at 3 DPI (22). While astrocyte reactivity was not noticeable on its own in the current study, ongoing studies are assessing fibrinogen extravasation and astrocytic involvement in pigs at these chronic time points.

Finally, we conducted histological analyses to assess the corpus callosum following injury. No overt APP pathology was detected in the corpus callosum and there were no significant changes in skeletal analysis features compared to sham. In addition to skeletal analysis, we also measured the thickness of the corpus callosum and the staining intensity of LFB to measure potential demyelination; however, there were no significant changes. Johnson et al. had previously found that the thickness of the corpus callosum was reduced with survival greater than 1 YPI in moderate-to-severe TBI patients, and that ongoing axonal degeneration was paired with CD68+ microglia (20). Indeed, progressive white matter pathology and corpus callosum volume loss has been noted in both human TBI and other animal models of TBI (5,6,50). Yet in this closed-head rotational model of single mild TBI, no significant changes to the corpus callosum have been detected out to 1 YPI. Examining specimens for corpus callosum deficits at time points greater than 1 YPI may be difficult to conduct, but they may be necessary to truly assess any potential chronic degeneration. Additional immunohistochemical staining, such as myelin basic protein, and genomic profiling may yield other pathological details into the state of the corpus callosum over time.

In summary, we demonstrated that a single, closed-head, mild TBI is associated with axonal pathology out to 7 DPI and alterations to microglia homeostasis out to chronic time points in certain anatomical subregions. This suggests that TBI can produce persistent changes to the neuroimmune response. Microglia activity has profound clinical significance as microglia-related pathways have become increasingly linked to Alzheimer’s disease and other neurodegenerative disease pathogenesis (58). Moreover, microglia may contribute to synapse loss, which is correlated to cognitive decline (49). However, further experimentation is needed to supplement our histopathology. Changes to gene expression followed by *in situ* hybridization may provide a wide array of inflammatory markers within an appropriate spatial context. Additionally, characterization of complement system activation and cytokine production may identify therapeutic targets. Finally, MRI or other neuroimaging studies would allow us to track neuroinflammation and white matter degeneration in a single specimen over time, providing us additional details that terminal time point pathology cannot provide. While it is difficult to obtain and characterize human tissue for corresponding analysis, it is hoped that further understanding of the chronic neuroinflammatory sequela of mild TBI in preclinical models such as this will translate to advances in the treatment and prevention of the long-term consequences of TBI in people.

## Acknowledgements

The authors thank Dr. John O’Donnell, Dr. Patricia Shewokis, Dr. Victoria Johnson, and Cassidy Fetterman for technical assistance.

## Author Contributions

MRG, JED, JAW, and DKC designed the experiments; MRG, DPB, NT, KLW, JPH, and KDB performed research; MRG and NP analyzed data; MRG and JED wrote the paper. All authors read and approved the final manuscript.

## Funding

This work was made possible through financial support provided by the Department of Veterans Affairs [RR&D Merit Review I01-RX001097 (Duda), BLR&D Merit Review I01-BX003748 (Cullen), and RR&D Career Development Award IK2-RX001479 (Wolf)], the National Institutes of Health [NINDS R03-NS116301 (Cullen), NINDS R01-NS101108 (Wolf), NINDS T32-NS043126 (Harris), NINDS F32-NS116205 (Wofford)], the CURE Foundation [Taking Flight Award (Wolf)], and the US Department of Defense [ERP CDMRP #W81XWH-16-1-0675 (Wolf)]. None of the funding sources aided in the collection, analysis, and interpretation of data, in the writing of the report, or in the decision to submit the paper for publication.

## Compliance with Ethical Standards

### Conflict of Interest

No competing financial interests exist. The authors have no conflicts of interest related to this work to disclose.

### Availability of Data and Materials

The datasets used during the current study are available from the corresponding author on reasonable request.

## Figure legends

**Supplementary Figure 1**. All Iba-1 positive cells were manually selected within the 40x region (a; scale = 20 μm). The image was deconvoluted, converted to binary, skeletonized, and skeletons not associated with manually selected cells were removed (b; scale = 20 μm). The Analyze Skeleton plugin on ImageJ was applied to all selected skeletons to quantify skeletal analysis features, such as branches (in red), junctions (arrow), and endpoints (arrowhead) (c).

**Supplementary Figure 2**. APP pathology burden at 230-270 rad/sec and 165-185 rad/sec, as well as non-linear best-fit lines for 7 DPI specimens, are shown for the fimbria/fornix (a-c), striatum (d-f), dorsal thalamus (g-i), ventral thalamus (j-l), and cerebellum (m-o). There was a significant increase in APP pathology in the fimbria/fornix at 7 DPI compared to sham at 230-270 rad/sec (p = 0.0072).

## Notes

### Competing Interest Statement

The authors have declared no competing interest.

