## Supplementary figures and images for "Diverse Changes in Microglia Morphology and Axonal Pathology Over One Year after Mild Traumatic Brain Injury in Pigs"

### Supplemental Figure 1

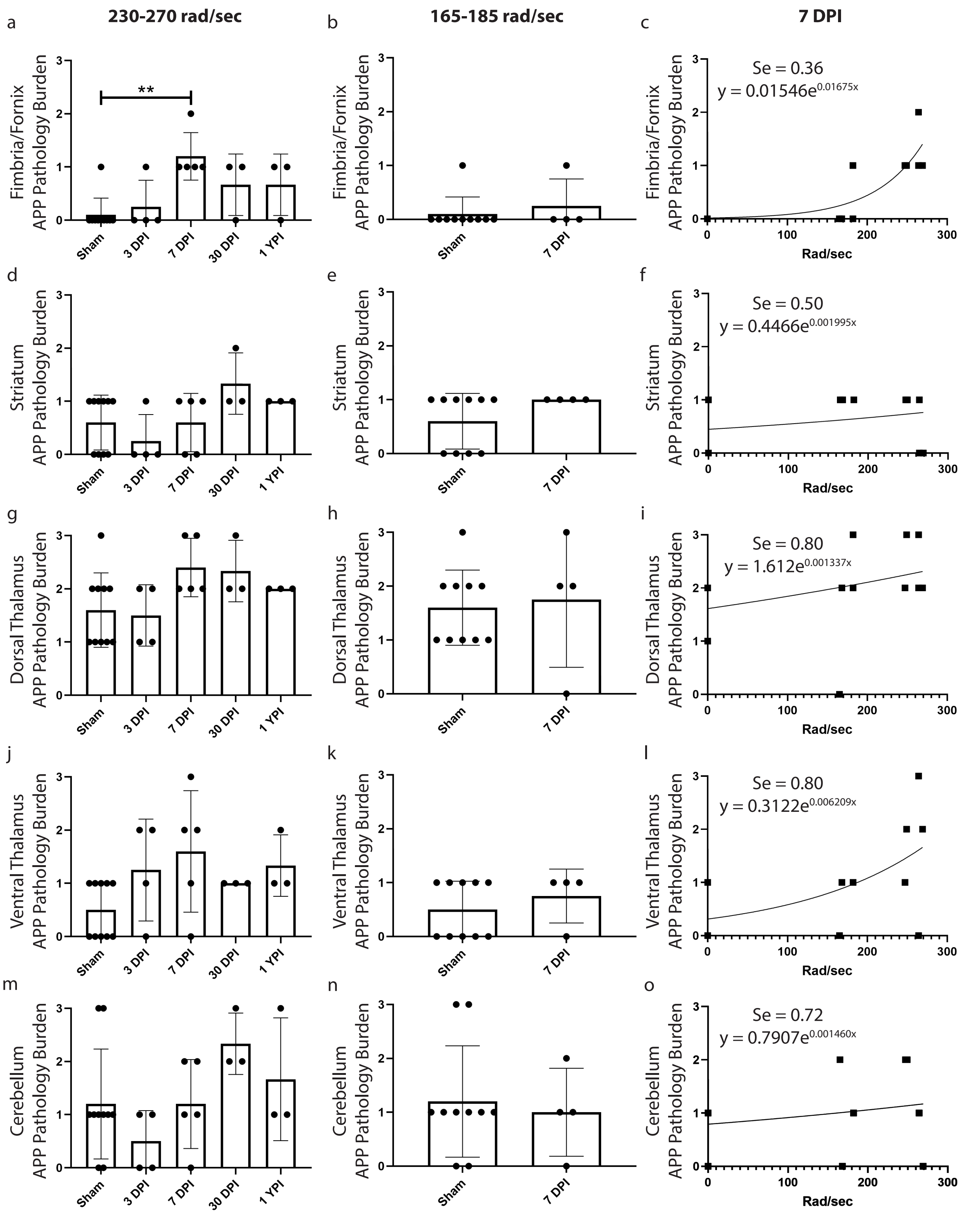

### Supplemental Figure 2

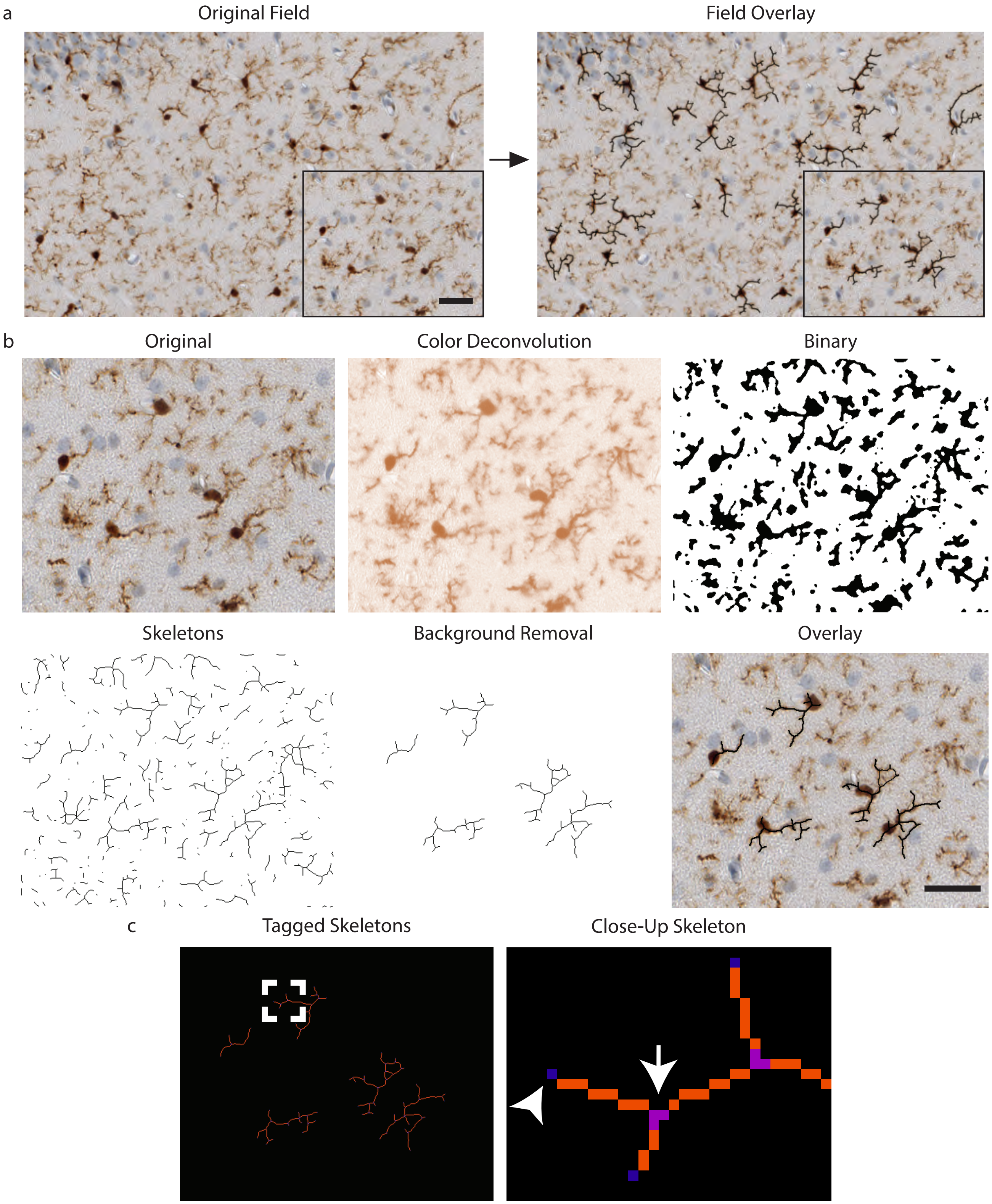
